# Establishing a reproducible approach for the controllable deposition and maintenance of plants cells with 3D bioprinting

**DOI:** 10.1101/2022.03.25.485804

**Authors:** Lisa Van den Broeck, Michael F Schwartz, Srikumar Krishnamoorthy, Ryan J Spurney, Maimouna Abderamane Tahir, Charles Melvin, Mariah Gobble, Rachel Peters, Atiyya Muhammad, Baochun Li, Maarten Stuiver, Timothy Horn, Rosangela Sozzani

## Abstract

Capturing cell-to-cell and cell-to-environment signals in a defined 3 dimensional (3D) microenvironment is key to study cellular functions, including cellular reprogramming towards tissue regeneration. A major challenge in current culturing methods is that these methods cannot accurately capture this multicellular 3D microenvironment. In this study, we established the framework of 3D bioprinting with plant cells to study cell viability, cell division, and cell identity. We established long-term cell viability for bioprinted *Arabidopsis* root cells and soybean meristematic cells. To analyze the large image datasets generated during these long-term viability studies, we developed an open source high-throughput image analysis pipeline. Furthermore, we showed the cell cycle re-entry of the isolated *Arabidopsis* and soybean cells leading to the formation of microcalli. Finally, we showed that the identity of isolated cells of *Arabidopsis* roots expressing endodermal markers maintained longer periods of time. The framework established in this study paves the way for a general use of 3D bioprinting for studying cellular reprogramming and cell cycle re-entry towards tissue regeneration.

## Introduction

A tight coordination between cellular reprogramming and cell cycle progression via cell-to-cell communication is required for tissue regeneration. In plants and animals, cell-to-cell communication and cellular reprogramming are both influenced by mechanical and positional cues (1). Thus, recapitulating the multicellular three-dimensional (3D) microenvironment behind cell-to-cell and cell-to-environment communication is key to gaining insights into the mechanical and positional context that governs cellular reprogramming towards tissue regeneration.

Multiple studies have established that conventional cell maintenance methods do not entirely mimic the 3D microenvironment observed in natural biological conditions in mammalian cells (2). 3D bioprinting enables the generation of specifically designed 3D cellular constructs. 3D bioprinting can be performed through the application of one or multiple modalities for the precise spatial deposition of biologically active material, referred to as a bioink. Specifically, 3D bioprinting provides an amenable and controllable system to fabricate complex systems that capture cellular dynamics and interactions in a physiologically accurate manner towards tissue regeneration (3-6). For example, human induced pluripotent stem cells have been bioprinted to study cell fate, phenotypic variation, and tissue regeneration (4). Plants inherently show extreme regenerative capacities. However, the regenerative capacities of individual isolated plant cells are greatly affected by cell type, tissue type, genotype, and the specific 3D microenvironment (1). With 3D bioprinting the mechanical and positional cues can be controlled by depositing living plant cells in a well-defined environment. Here, we used *Arabidopsis* root-derived protoplasts as a model system to showcase the potential of 3D bioprinting to study cell viability, cell division and cellular reprogramming in a tunable microenvironment. Specifically, we maintained bioprinted *Arabidopsis* root cells isolated from two different tissue types for up to 7 days. In this timeframe, we showed that the percentage of isolated cells expressing endodermal markers increases, indicating that the identity of isolated cells changes over longer periods of time. Lastly, we demonstrated the applicability of 3D bioprinting for studying regeneration of isolated single cells for model and crop species.

## Results

### Demonstrating the maintenance of 3D bioprinted *Arabidopsis* root cells

Cellular functions such as cell differentiation, cell cycle progression, responsiveness to stimuli, and cellular reprogramming are driven by cell-to-cell and cell-to-environment communication. These cell-to-cell and cell-to-environment signals are not entirely captured in suspension cell cultures due to lack of a natural microenvironment (7-9). On the other hand, 3D encapsulation methods can replicate the natural microenvironment. Moreover, they can immobilize plant cells allowing for more physiologically accurate cellular behavior. We utilized 3D bioprinting to recapitulate the plant cell’s microenvironment and behavior. For this, we optimized two key aspects of successful bioprinting, which includes the bioink and bioprinting parameters. The bioink is a combination of living cells and the scaffold material that provides support to the cells and the nutrients needed to ensure cell viability (10). To achieve long-term cell viability, we optimized tissue-specific bioinks and established a reproducible experimental pipeline that consists of bioprinting protoplasts, and maintaining and imaging the bioprinted constructs over time (Fig 1A). In this pipeline, we focused on *Arabidopsis thaliana* roots, a tissue of which the maintenance outside its context is known to be technically challenging (11). Additionally, to further gain insights into whether different tissue types would be more prone to cellular reprogramming, we isolated protoplasts from both meristematic and differentiated root tissue and performed comparative analyses. These comparative analyses and time course experiments resulted in large-scale image datasets to process and analyze. To analyze these large image datasets generated from our bioprinting pipeline, we developed a semi-automated and high-throughput image analysis pipeline enabling the quantification of cells in confocal microscopy z-stack images (Methods). To make it freely available for the scientific community, the entire pipeline was wrapped in a graphical user interface (Fig 1B).

**Fig. 1.**
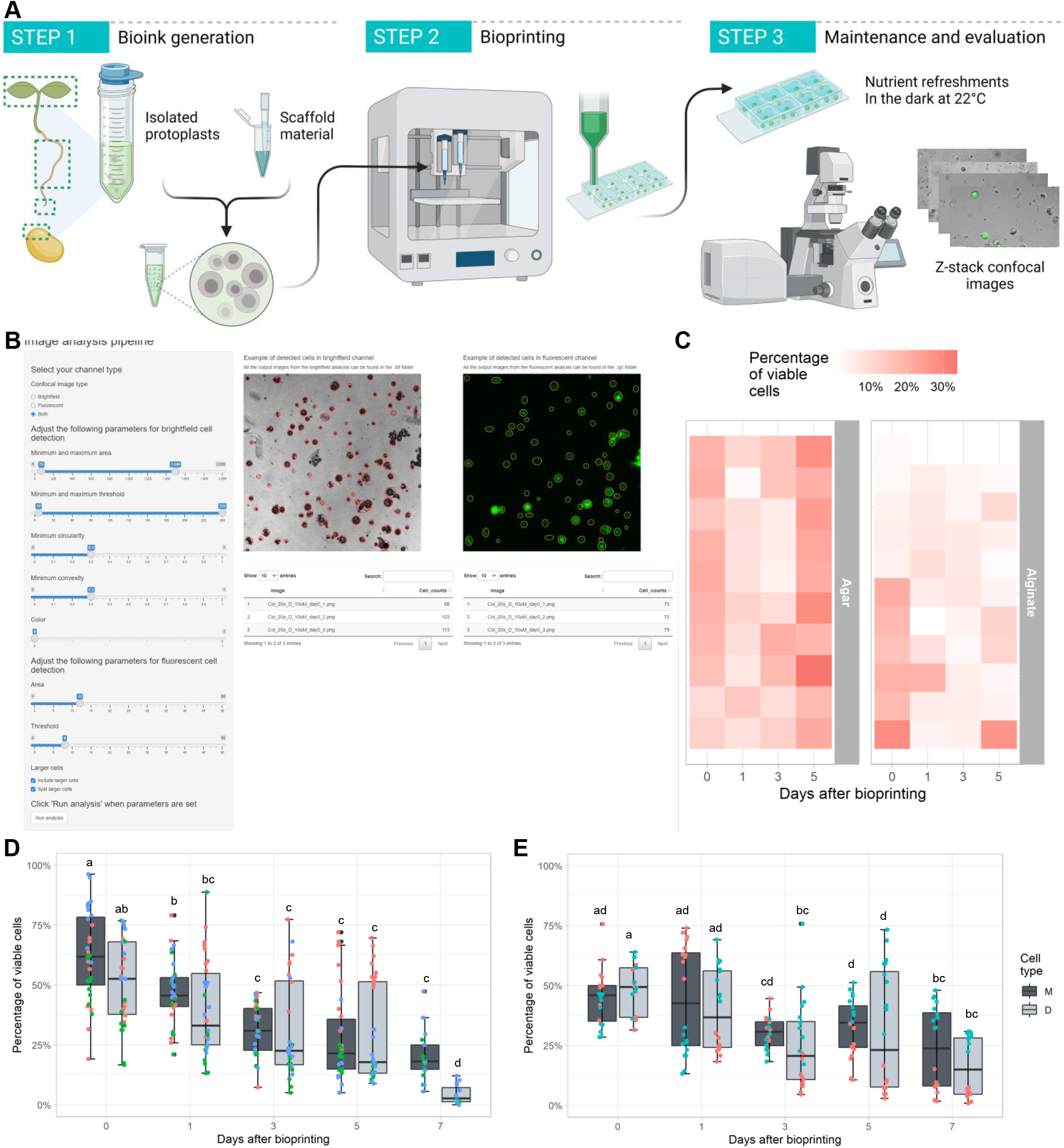
Maintenance of bioprinted cells isolated from Arabidopsis meristematic root tissue. (A) Schematic overview of the key steps in our 3D bioprinting protocol (created with BioRender.com). (Step 1) Bioink is generated by mixing root protoplasts and scaffolding material, (Step 2) 3D bioprinting with the CELLINK BIOX, and (Step 3) maintaining and imaging over time. (B) Screenshot of the graphical user interface of our in-house developed pipeline enabling the high-throughput quantification of cells in confocal z-stack images. (C) The percentage of viable cells in different bioprinted cellular constructs over time using two different scaffolding materials; 0.6% low melting agar and 0.75% sodium alginate. (D-E) The percentage of viable cells isolated from Col-0 meristematic (M) and differentiated (D) root tissue upon bioprinting (D) or when deposited manually using a micropipette (E). Cell viability was evaluated with fluorescein diacetate (FDA) staining immediately after bioprinting (day 0) and 1, 3, 5 and, 7 days after bioprinting. Colored dots on box plots represent the independent experiments (B), number of bioprinted constructs (N) = 30 (D), N = 20 (E), N = 10 (C), letters indicate significant differences (p < 0.05, Tukey test).

Tissue-specific bioinks require the balancing of auxin and cytokinin, which is critical for long term cell viability and proliferation (1). Additionally, several other compounds have been identified to activate the intrinsic cellular programs that trigger cell proliferation. To establish a reproducible experimental pipeline using plant cells for 3D bioprinting that supports viable cells for extended periods, we used optimized concentrations of auxin and cytokinin, and added phytosulfokine (PSK) and folic acid, two compounds that stimulate cell proliferation, to the bioink (12-14) (see Methods). Additionally, to avoid the formation of a thermal gradient within the bioink and our cells, we incorporated an axial piston in the cartridge. After optimization, a usual experiment yielded about 62.2%±8.2% and 51.6%±8.3% viable meristematic and differentiated protoplasts immediately after 3D bioprinting, and less of the meristematic and differentiated isolated cells (27.7%±7.6% and 29.5%±3.5%) were generally still viable after 5 days (Fig 1D, Supplemental Fig S1A-B, Supplemental Data S1). Given that after bioprinting 40% of the *Arabidopsis* root protoplasts were not viable upon FDA staining, we compared the viability of bioprinted cells to manually pipetted cells. Manually pipetted meristematic and differentiated protoplasts showed a slightly reduced but comparable viability to bioprinted cells of 44.9%±6.6% and 48.6%±6.6%, respectively, indicating that the process of bioprinting has no observable detrimental effect on cell viability but rather the protoplast isolation (Fig 1E, Supplemental Data S1).

3D bioprinting requires a suitable scaffold that is biocompatible, ensures transfer of nutrients and oxygen to the cells, and offers structural support (15). In our experimental pipeline, we chose low melting agarose as scaffold due to its established biocompatibility and reversible thermal crosslinking properties resulting in a more tunable operational workflow (15). However, it was previously reported that *Arabidopsis* shoot cells encapsulated in sodium alginate were viable and able to regenerate (16,17). To test the compatibility of sodium alginate with bioprinting *Arabidopsis* root cells for prolonged viability, we bioprinted protoplasts in the two different scaffold materials with different properties, namely 0.6% (w/v) low melting agarose and 0.75% (w/v) sodium alginate (16,17). As expected, after 5 days, we observed 23.6%±5.8% viability within the agarose compared to 7.7%±1.3% with the sodium alginate (Fig 1C, Supplemental Data S1), and after 7 days, we observed mechanical deformations in the alginate constructs in some cases (Supplemental Fig S1C). We reasoned that this was due to alginate requiring ionic crosslinking by the addition of bivalent cations, such as calcium ions, which adds complexity to the 3D bioprinting process. Therefore, we chose to primarily utilize low melting agarose as the scaffold material in this study.

Overall, we demonstrated the utilization of extrusion-based 3D bioprinting to reliably deposit plant cells isolated from different tissue types and demonstrated the possibility of using different scaffold materials in the bioprinting process. Importantly, we showed the maintenance of *Arabidopsis* root protoplasts within 3D bioprinted constructs over prolonged periods of time while maintaining sufficient viability.

### Applicability of 3D bioprinting towards plant cell regeneration

Bioprinting isolated cells into designed cellular constructs provides a system to evaluate cellular regeneration. Previous studies have shown that cellular regeneration by callus formation from root explants initiates with divisions of pericycle cells, rather than dedifferentiation and cell cycle re-entry of root cells (17). To our knowledge, cell cycle re-entry of protoplasts derived from *Arabidopsis* roots has not yet been achieved (11,17). To understand whether bioprinted meristematic and differentiated root cells enter the cell cycle, we evaluated the bioprinted cells for divisions and microcallus formation. We observed that bioprinted *Arabidopsis* root cells undergo their first cell division as early as 5 days (Fig 2A). To map the potential of cells to re-enter the cell cycle upon bioprinting, we used the cell cycle marker pCYCB1;1:CYCB1;1-GFP, which is expressed at the transition from the G2 gap phase into mitosis (M-phase) (18). Over the evaluated time course, on average ∼5% and ∼2% of the bioprinted meristematic and differentiated root cells expressed CYCB1;1, respectively, indicating the restricted potential of protoplasts to re-enter the cell cycle (Fig 2B, Supplemental Data S1). After a 7 to 10 day period, ∼12% of the bioprinted constructs developed a microcallus, here defined as 4 cells or more (Supplemental Data S2). These results suggest that under these conditions the bioprinted *Arabidopsis* root cells have the ability to undergo cell divisions and form microcalli.

**Fig. 2.**
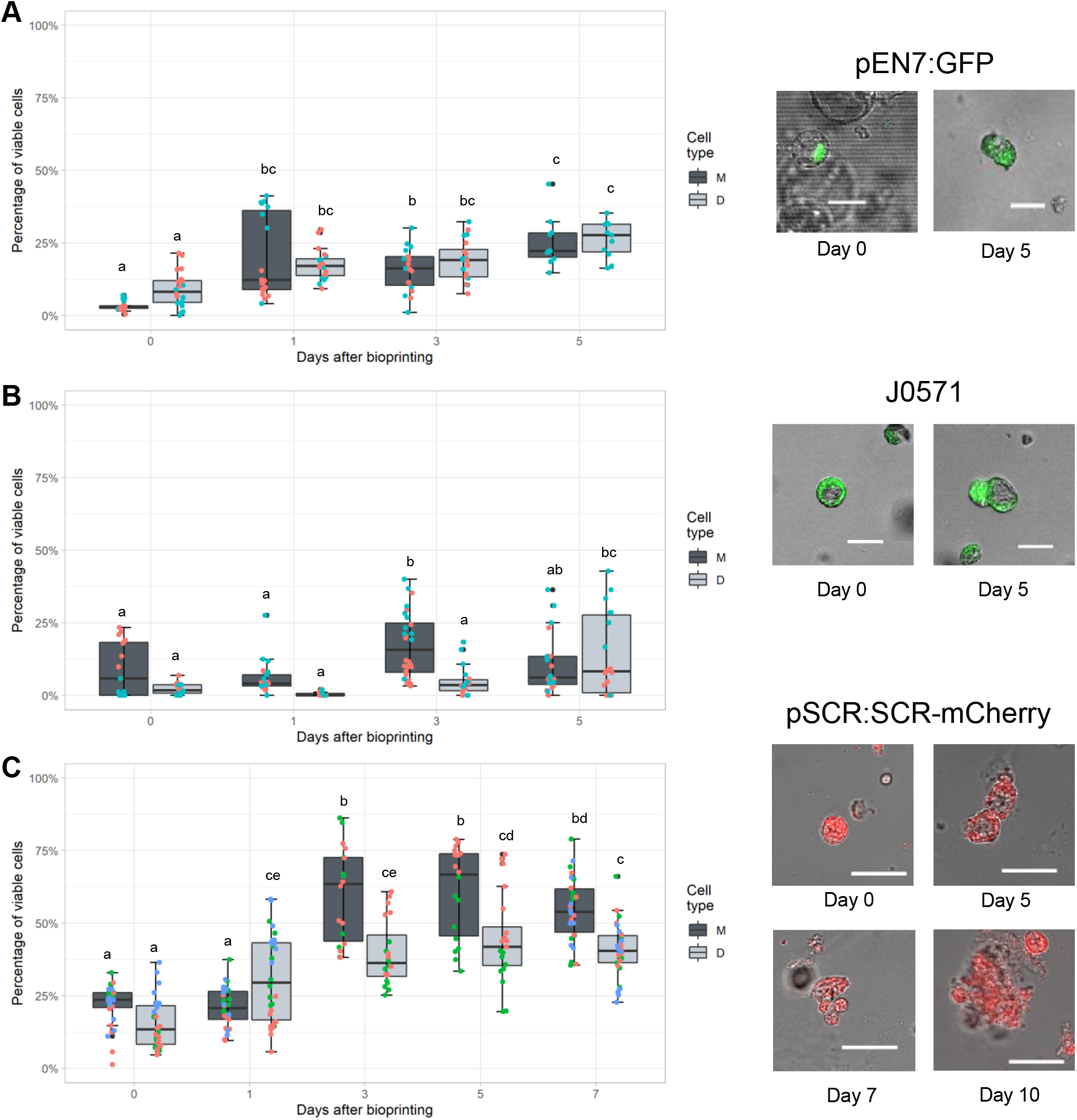
Isolated cells from Arabidopsis roots, shoots, and shoot meristematic region of soybean embryonic axis divide and form microcalli. (A) From left to right: high resolution confocal microscopy images of an isolated protoplast, an isolated protoplast that has undergone cell division, and microcallus formed from a protoplast isolated from Arabidopsis roots, shoots or soybean shoot meristematic region. (B) Protoplasts were isolated from pCYCB1;1:CYCB1;1-GFP meristematic (M) and differentiated (D) root cells. The cells of the same 3D bioprinted structure were imaged immediately after bioprinting (day 0) and 1, 3, 5, and 7 days after bioprinting. The protoplasts were bioprinted in protoplast induction media with 0.6% low melting agar. Colored dots represent the independent experiments, Number of bioprinted constructs (N) = 30, letters indicate significant differences (p < 0.05, Tukey test).

The division, maintenance, and regeneration of *Arabidopsis* shoot protoplasts is well established in prior studies (17,19). To compare our current system with prior data, we bioprinted shoot protoplasts using our bioink optimized for root-derived cells. Similar to root cells, bioprinted shoot cells showed an initial viability of 57.6%±4.3% (Supplemental Fig 2A-B). After 5 days the viability was reduced to 20.5%±2.2% and was observed to be further diminished after 7 days (Supplemental Fig 2A). Although we observed cell divisions as early as 5 days, we did not detect any microcallus formation within the imaged timeline. We speculate that the maintenance of cells derived from different tissue types or species need the utilization of bioink and scaffold materials that are specifically optimized. Thus a specially optimized bioink might be necessary stimuli for microcallus formation from shoot tissue. Similarly, we reasoned that successful 3D bioprinting requires optimized bioinks for each tissue type and further optimization is necessary across species. To demonstrate the extension of bioprinting to crops for studying regeneration, we first optimized a bioink for soybean protoplasts isolated from embryonic shoot meristems (see methods) (20-27). To this end, we tailored the bioprinting parameters due to the challenges associated with bioprinting soybean protoplasts. Because the protoplast isolation utilizes a relatively short incubation time in the cell wall digesting enzymes, some tissue was partially digested. To avoid non-uniform extrusion due to nozzle blockages, we used a ∼ 2 times wider needle for bioprinting, which also had the added benefit of reducing shear stresses on the cells. Using this optimized pipeline, we bioprinted meristematic cells isolated from *Glycine max* (L.) Merr. cv Thorne (28) and evaluated cell viability, cell division, and microcallus formation (29) (Supplemental Fig 2D). Overall, isolated soybean cells showed 28%±9.6% viability immediately after bioprinting and 43%±11.2% viability after 14 days (Supplemental Fig 2D). We observed that some cells anisotropically elongate, which correlates with the onset of cell division as early as 3 days after bioprinting. Furthermore, some of these cells grow into an unorganized cell mass. After 14 days, we observed the formation of on average 5 to 6 viable microcalli in 90% of the bioprinted constructs (Fig 2A, Supplemental Data S2). These results indicate that bioprinting can be used as an enabling technology to study cellular regeneration in crops.

### Identity maintenance of 3D bioprinting root cells

The root contains multiple cell types and thus the bioprinted root cell population is highly heterogeneous. To shed light on the identity of the isolated cells that remain competent for cellular reprogramming leading to regeneration, we examined the cellular identity of our bioprinted root cells and the formed microcalli. To this end, we focused on the ground tissue identity and tracked it using cell-type specific marker lines. Specifically, we used a nuclear-localized transcriptional marker line for the endodermis and cortex endodermis initial (CEI) cells, pEN7:GFP, the translational fusion pSCR:SCR-mCherry to identify the endodermis, CEI, and QC, and the enhancer trap J0571 to mark the endodermis and cortex (Supplemental Fig 3) (30). Immediately after bioprinting we observed between 5.4%±1.6% and 10.1%±2.2% of meristematic and differentiated protoplasts with EN7 expression, which is comparable to the fractions of endodermal cells in an intact root (Supplemental Table S1). The percentage of EN7-expressing cells increases over time to 25.3%±8.7% and 26.3%±6.3% at 5 days for meristematic and differentiated root cells, respectively (Fig 3A). We observed a similar trend with the translational fusion SCR:SCR-mCherry where the marker’s expression increases in both meristematic and differentiated tissues to 54.4%±6.5% and 40.7%±4.1% at 7 days, respectively (Fig 3C). Thus, for both endodermal markers, we could observe an increase in cells expressing the markers over time, which is in line with previous studies that show that isolating protoplasts induces enhanced expression variation at the genome level and promotes stochastic activation of gene expression (19). Similarly, previously published RNAseq and ATACseq data of leaf isolated protoplasts showed an induction of EN7 and SCR expression and chromatin accessibility upon protoplast isolation (19) (Supplemental Table S1). Interestingly, for pEN7:GFP, both meristematic and differentiated bioprinted cells showed an equal fraction of fluorescent cells, while SCR expression was significantly increased in meristematic root cells compared to differentiated root cells (Fig 3). Additionally, we analyzed the enhancer trap line J0571, which includes the additional fluorescence of the cortex (30). Similar to pEN7:GFP, J0571-derived protoplasts showed no prominent differences between meristematic and differentiated bioprinted cells (Fig 3B). Conversely, the percentage of GFP-expressing cells from J0571 did not increase over time. Overall, the changes in the percentage of cells expressing the endodermal markers indicate that the identity of isolated cells changes over longer periods of time, which could be the result of a stochastic increase in ectopic expression or the loss of non-cell autonomous signals.

**Fig. 3.**
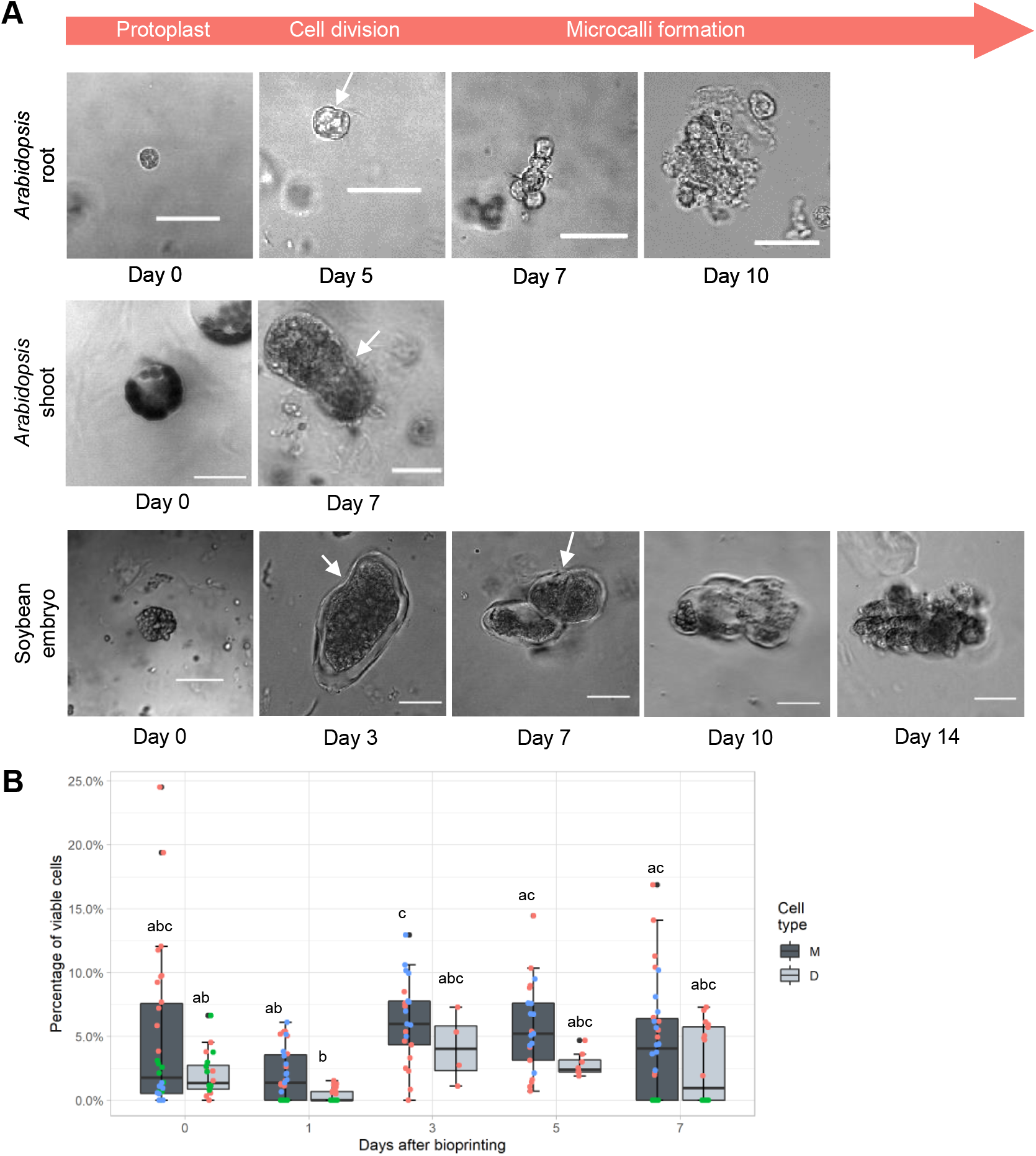
Ground tissue identity of isolated root cells upon bioprinting. (A-C) Protoplasts were isolated from pEN7:GFP (A), J0571 (B), and pSCR:SCR-mCherry (C) meristematic and differentiated root cells. The cells of the same 3D bioprinted structure were imaged immediately after bioprinting (day 0) and 1, 3, 5, and 7 days after bioprinting. Representative high resolution confocal images of 3D bioprinted cells are given. Colored dots represent the independent experiments, Number of bioprinted constructs (N) = 20 (A, B), N = 30 (C), * = p < 0.05 (Tukey test).

To further gain understanding of the identity of the microcalli, we evaluated whether the formed microcalli showed expression of our marker lines. As such, we could at least partially identify which lineage-specific cells were reprogrammed to pluripotency. Because regenerating cells are very sparse, it is challenging to investigate this process. We only identified sufficient microcalli in the bioprinted constructs of pSCR:SCR-mCherry. Namely, 100% of the pSCR:SCR-mCherry microcalli (7 in total) expressed mCherry fluorescence, which suggested that endodermal, CEI, and/or QC-derived cells were capable of proliferating. Additionally, we identified microcalli expressing the pCYCB1;1:CYCB1;1-GFP reporter. Of the pCYCB1;1:CYCB1;1-GFP microcalli, 33% showed GFP fluorescence, which suggests that 3D bioprinted cells are actively dividing.

## Discussion

Isolated single cells provide a great system for studying aspects of cell physiology, cell identity, cell-to-cell communication, and cell cycle re-entry. Here, we were able to maintain isolated *Arabidopsis* root cells with high viability for multiple days using a reproducible 3D bioprinting pipeline. Our bioprinting pipeline and semi-automated image analysis pipeline allowed us to evaluate isolated *Arabidopsis* cells, including several marker lines, through their developmental lineage. The percentage of isolated cells expressing endodermal markers increases over time, suggesting that the bioprinted cells might be either partially changing identity or transitioning to a previously unknown identity. A previous study in *Arabidopsis* leaf protoplasts suggests that protoplasts undergo increased stochastic and ectopic gene expression multiple days after isolation (19). Thus an increased ectopic expression might result in the observed changes in expression of identity genes. Another hypothesis is that the loss of an organizational structure, like that of an intact root, leads to non-cell autonomous signals reaching otherwise unreachable cells or not reaching their intended neighboring cells. This loss and gain of molecular signals and the following disruption in their respective signaling pathways could result in cell identity changes. To further gain insights into these two hypotheses, it would be essential to develop single cell RNAsequencing protocols compatible with encapsulated cells.

Our pipeline allows us to design the cell’s local environment to recreate conditions that sustain cell viability and microcallus formation (Fig 2). Cells isolated from meristematic and differentiated *Arabidopsis* root tissue were able to re-enter the cell cycle and form microcalli, indicating a key role for the 3D environment. In line with this hypothesis, preliminary studies in which transgenic *Arabidopsis thaliana* were cultured within tailored 3D fibrous microenvironments were observed to show growth and morphological characteristics of greater complexity than those cultured on standard 2D scaffolds (31). Interestingly, the formed microcalli expressed the endodermal marker SCR. Previous studies showed that callus formation from roots explants was initiated by pericycle cells differentiating into cells resembling root meristems (11,17). However, the expression of SCR in the bioprinted microcalli indicates that in addition to pericycle-derived microcallus formation, another reprogramming mechanism might trigger microcallus formation. The underlying molecular basis of microcallus formation from a single cell are of interest to enhance regeneration of recalcitrant species, increase transformation events, and enable gene editing studies. In this study, we applied our bioprinting pipeline to reprogram meristematic soybean cells to pluripotency. We isolated cells from soybean embryonic axes and were able to show the formation of microcalli after 14 days, a comparable timeline with previous studies (32). Further identifying structural polymers, signaling molecules, or cell types that generate cell-type tailored environments to ensure efficient plant regeneration could have a large number of applications.

Our study provides the groundwork for utilizing 3D bioprinting in the plant sciences field and details protocols and guidelines. However, more research needs to be done. Specifically, the identification and detailed classification of the dimension parameters and signaling factors that stimulate plant cell processes, such as proliferation, cell attachment, regeneration, and differentiation, are crucial. For example, identifying compounds or cell types that ensure efficient plant regeneration or other cellular responses could be of interest in a wide range of biotechnological applications. In the future, scaffolds and bioinks based on a variety of synthetic and naturally occurring substrates will need to be designed in a case-by-case manner to engineer specific plant cell-based models and support 3D growth of cells. Additionally, external stimuli to induce time-dependent changes in functionality of encapsulated cells may be adopted. Overall, we have shown the potential of 3D bioprinting to study cell viability, cell division and cellular reprogramming in a tunable microenvironment.

## Methods

### Plant lines and growth conditions

The following lines were used: wild type, pEN7:GFP (AT4G28100) (33), pSCR:SCR-mCherry (AT3G54220) (34), and pCYCB1;1:CYCB1;1-GFP (AT4G37490) (35) in Columbia-0 background, and J0571 (30) in the C24 background. *Arabidopsis thaliana* seeds were surface sterilized using 50% bleach and 0.05% Tween-20 for 5 minutes, followed by a 2 minute incubation in 70% ethanol for 2 minutes. Seeds were rinsed with sterile deionized water at least 7 times. Following the last rinse, the seeds were stratified at 4°C for 2 days and plated on 1× Murashige and Skoog (MS) agar supplemented with sucrose (1% sucrose total) on top of Nitex mesh [Genesee]. All plants were grown in a vertical position at 22°C in long-day conditions (16-h light/8-h dark cycle).

Soybean (*Glycine max* (L.) Merr. cv Thorne) seeds were surface sterilized using 70% ethanol for 2 minutes, followed by a 10 minute incubation in 50% bleach and 0.05% Tween-20. Seeds were rinsed in sterile deionized water 10 times. Following the last rinse, the seeds were placed in a sterile petri dish, covered with sterile deionized water, and imbibed at 27°C in long-day conditions (16-h light/8-h dark cycle) for 24 hours.

### Isolation of protoplasts

To avoid contamination, all steps were performed in a laminar flow hood. To prepare the enzyme solution, 0.45 g cellulase [EMD Millipore] and 0.03 g pectolyase [Sigma-Aldrich] were dissolved in a 30 mL solution containing: 5.465 g of mannitol, 0.05 g of 0.01% BSA, 500 μL 0.2 M Magnesium chloride, 500 μL 0.2 M calcium chloride, 500 μL 1 M MES, 500 μL 1 M potassium chloride, 50 mL deionized water, and pH was set to 5.5 with Tris-HCL. The enzyme was sterilized using a 0.20 μm syringe filter. For each sample, 7 mL of fresh enzyme solution was pipetted into 35-mm-diameter petri dishes. We avoided creating bubbles when pipetting the enzyme solution to avoid cell lysis at later stages in the protocol.

A 70 µm cell strainer was placed in each 35-mm-diameter petri dish. Approximately 1-2 mm of the root tip was cut to isolate the meristematic region of the root and put into the strainer in enzyme solution. The samples were incubated for 2 hours at 85 rpm at room temperature with supplemental stirring every 30 minutes. Next, all of the cells and enzyme solution were transferred to a 15 mL conical tube and centrifuged for 6 min at 200 g. The supernatant was removed and the pellet was resuspended with 1 mL protoplast induction media (PIM) (36,37). PIM (*Arabidopsis*) contains a cocktail of growth hormones, sugar, mannitol to ensure a proper osmolarity, folic acid (13,14), phytosulfokine (PSK) (12,40). The resuspended solution was transferred to a 70 μm filter placed on top of a 50 mL conical tube.. All the filtered liquid was transferred to a 40 μm filter placed on top of a 50 mL conical tube. The subsequent filtered liquid contains the protoplasts used for bioprinting. The volume of cells in each solution was estimated using a hemocytometer. The starting cell densities are listed in Table 2.

For soybean protoplast isolation, 2% (w/v)cellulase [EMD Millipore] and 0.03% (w/v) pectolyase [Sigma-Aldrich] were dissolved in a 30 mL solution containing: 13% mannitol, 27.2mg/l KH_2_PO_4_, 100 mg/l KNO_3_, 150 mg/l calcium chloride, 250 mg/l magnesium chloride, 2.5mg/l Iron (III) sulfate, 0.16 mg/l KI, 50 mL deionized water, and pH was set to 5.5 with Tris-HCL. The enzyme was sterilized using a 0.20 μm syringe filter. For each sample, 7 mL of fresh enzyme solution was pipetted into 35-mm-diameter petri dishes. We avoided creating bubbles when pipetting the enzyme solution to avoid cell lysis at later stages in the protocol.

A 70 µm cell strainer was placed in each 35-mm-diameter petri dish. Meristematic shoot tissue from embryonic axis from imbibed soybean seeds was isolated and put into the strainer in enzyme solution. The samples were incubated for 2 hours at 85 rpm at room temperature with supplemental stirring every 30 minutes. Next, all of the cells and enzyme solution were transferred to a 15 mL conical tube and centrifuged for 6 min at 200 g. The supernatant was removed and the pellet was resuspended with 1 mL bioink (Table 1). The resuspended solution was transferred to a 70 μm filter placed on top of a 50 mL conical tube. All the filtered liquid was transferred to a 40 μm filter placed on top of a 50 mL conical tube. The subsequent filtered liquid contains the protoplasts used for bioprinting. The volume of cells in each solution was estimated using a hemocytometer.

**Table 1.**
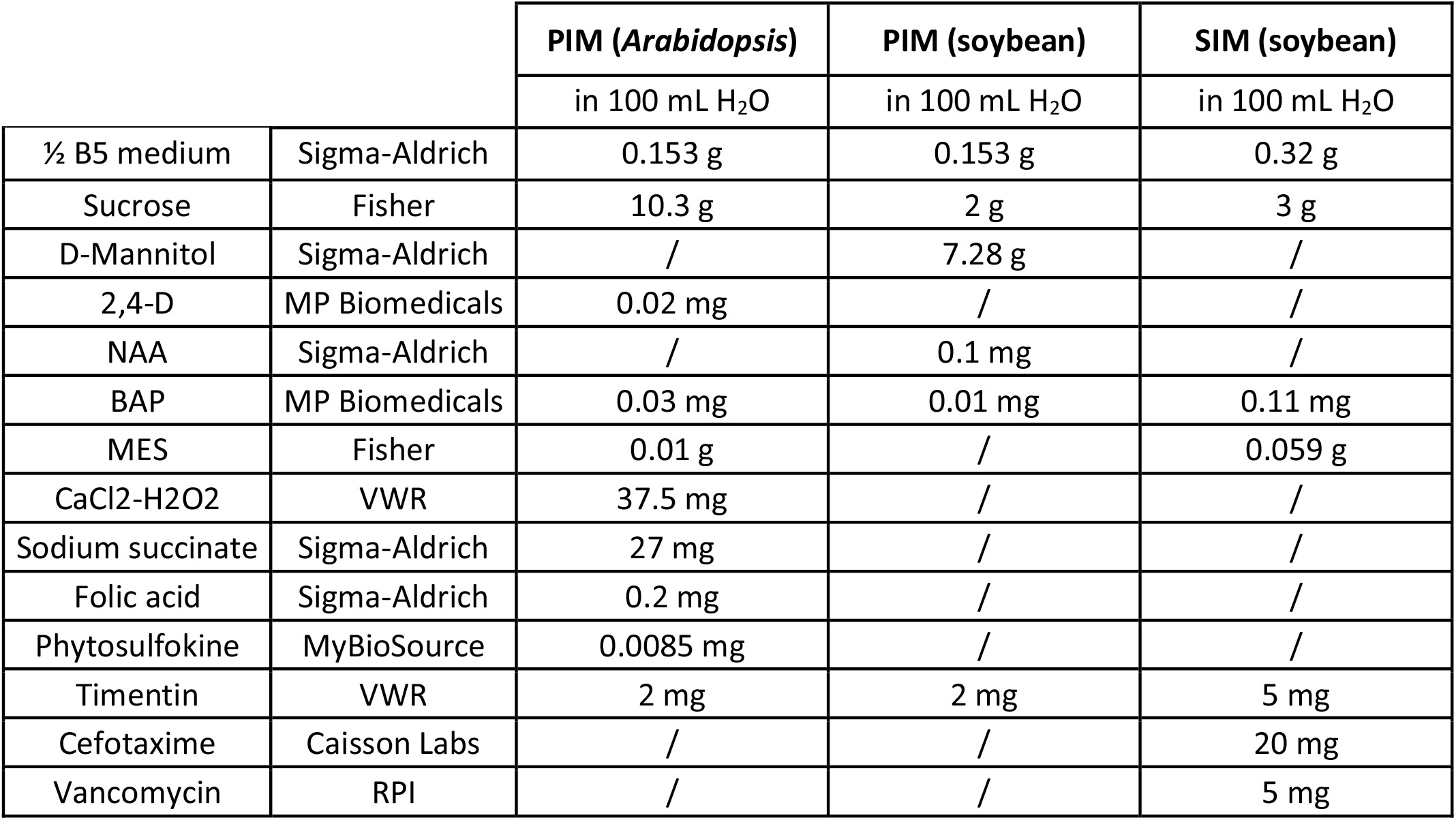
Recipe for PIM and SIM. The pH was set to 5.7 followed by autoclaving. Antibiotics were filter-sterilized and added after autoclaving. PIM = protoplast induction media, SIM = shoot induction media

### Bioink and 3D-bioprinter

The isolated protoplasts were resuspended in a specialized bioink that consisted of the PIM that was supplemented with 0.6% low melting agarose (Sigma-Aldrich) to provide support for cell growth. Namely, the PIM containing the protoplasts were mixed with a stock scaffold solution consisting of 2.4% Agarose dissolved in PIM, at a ratio of 3:1, to yield the desired bioink used for bioprinting.

The protoplasts were printed with a 3 mL syringe barrel that incorporated an axial piston, using an extrusion 3D bioprinter (CELLINK BIOX), into an 8-well µ-slide (IBIDI), which facilitated the acquisition of cell images at multiple time points. The bioink-embedded protoplasts are printed at a temperature of 37 degrees rendering liquid low melting agar. To minimize the effects of shear stress on the suspended cells during the 3D bioprinting process, the sterile blunt needle tip inner diameter and extrusion pressure were optimized. The optimized bioprinting parameters used are listed in Table 2. The extrusion of the protoplast-embedded bioink was followed by reversible thermal gelation of the agar by free convection, which results in a temperature reduction and solidification of the scaffold material, thus ensuring the fabrication of the desired 3D construct. The protoplast embedded bioink was extruded into 8-well µ-slides, with each well holding 4 bioprinted droplets. To prevent drying of the 3D constructs and allow for nutrient and hormone exchange, the cells are covered with 250 μL PIM in each well. Re-supplantation of the PIM was performed every 2 days. The bioprinted constructs were kept in the dark. For soybean bioprinted constructs, 250 μL shoot induction media (SIM) was used as refreshment at 7 and at 14 days after bioprinting.

**Table 2.** 3D bioprinting parameters.

| Parameter | *Arabidopsis* | Soybean |
| --- | --- | --- |
| Inner diameter needle tip | 210 $\mu\text{m}$ (27 G) | 413 $\mu\text{m}$ (22 G) |
| Extrusion pressure | 20 kPa | 20 kPa |
| Printhead | Extrusion | Extrusion |
| Extrusion temperature | 37 $^{\circ}\text{C}$ | 37 $^{\circ}\text{C}$ |
| Extrusion time | 0.8 s | 0.8 s |
| Printbed temperature | 25 $^{\circ}\text{C}$ | 25 $^{\circ}\text{C}$ |
| Average cellular density | 1392 ± 152 cells/$\mu\text{L}$ | 1379 ± 288 cells/$\mu\text{L}$ |

### Staining and confocal imaging

Prior to image acquisition, cells were stained with 0.01% fluorescein diacetate (FDA, Sigma Aldrich), a cell-permeable esterase substrate that measures intracellular enzymatic activity. Image acquisition was performed on a Zeiss LSM 880 confocal microscope. For each independent experiment, 10 bioprinted constructs were imaged by obtaining z-stack images with a width of 5 to 20 µm. Each z-stack contained between 50-300 cells. The obtained microscopy images were sorted per day, and each folder of images corresponding to a specific day was analyzed separately with our in-house developed pipeline, in order to improve counting strategies.

### Automated image analysis pipeline

To facilitate repeatable, robust, and rapid analysis of a large number of confocal images of cells, we generated an automated open source image analysis pipeline that is based on a can be run through an R Shiny graphical user interface (GUI). The GUI runs three in-house developed python scripts in the backend. The first python script separates the fluorescence and brightfield channels, performs a z-projection of the brightfield channel using minimum intensity, and saves the projections as .png files. This function relies on the pyimageJ package (41). The second python script quantifies cells in the brightfield channel using the publicly available OpenCV Computer vision library for tracking and counting of cells that performed contrast limited adaptive histogram equalization on the brightfield images, and used an edge detection algorithm to detect and count cells (42). The third script separates fluorescence and brightfield channels, performs z-projection of the fluorescence channel using maximum intensity, and detects and quantifies cells in the fluorescent channels with ComDet v 0.5.3, an open source plugin for ImageJ (43). This function also relies on the pyimageJ package (41). The program code for the entire image analysis pipeline contained in Python and further developed into an R Shiny application (44) can be easily accessed, along with the usage instructions, from the Github repository at https://github.com/LisaVdB/Confocal-z-stack-cell-detection.

## Data availability

Scripts for our high-throughput and automatic image analysis are available at https://github.com/LisaVdB/Confocal-z-stack-cell-detection.

## Acknowledgements

We thank the Cellular and Molecular Imaging Facility (CMIF) at North Carolina State University, specifically Mariusz and Eva, for their imaging advice. We thank coffee, sharp blades, petri dishes, and functional cellulase. This work was funded by the National Science Foundation (NSF) (EAGER MCB #2039285) to TH and RS.

---

**Supplementary Figure S1.**
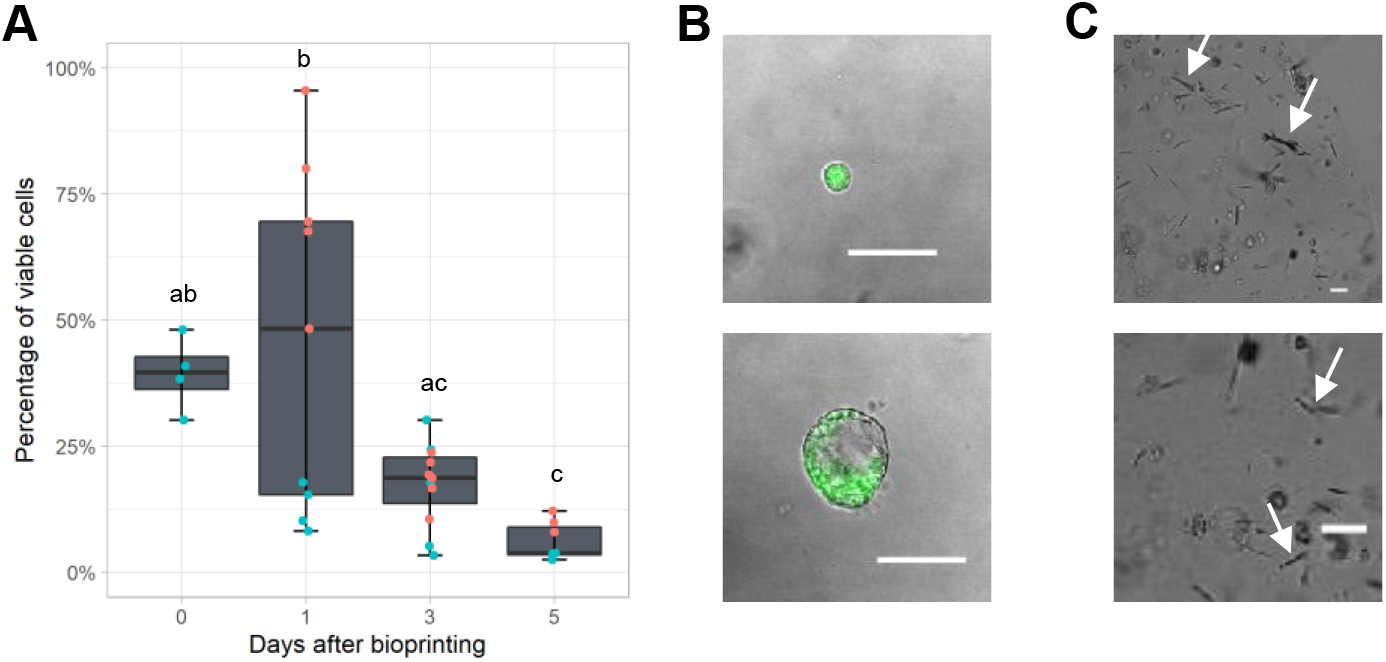
Maintenance of cells isolated from Arabidopsis meristematic root tissue. (A) Percentage of viable meristematic root cells when bioprinted with a pre-optimized set of parameters. Namely, direct pneumatic extrusion was utilized, and the media was not supplemented with PSK and Folic acid. (B) Representative high resolution confocal images of viable 3D bioprinted cells isolated from meristematic root tissue. (C) Bioprinted construct with alginate scaffolding. Arrows indicate mechanical deformations in the alginate constructs. Scale bar = 20 μm.

**Supplementary Figure S2.**
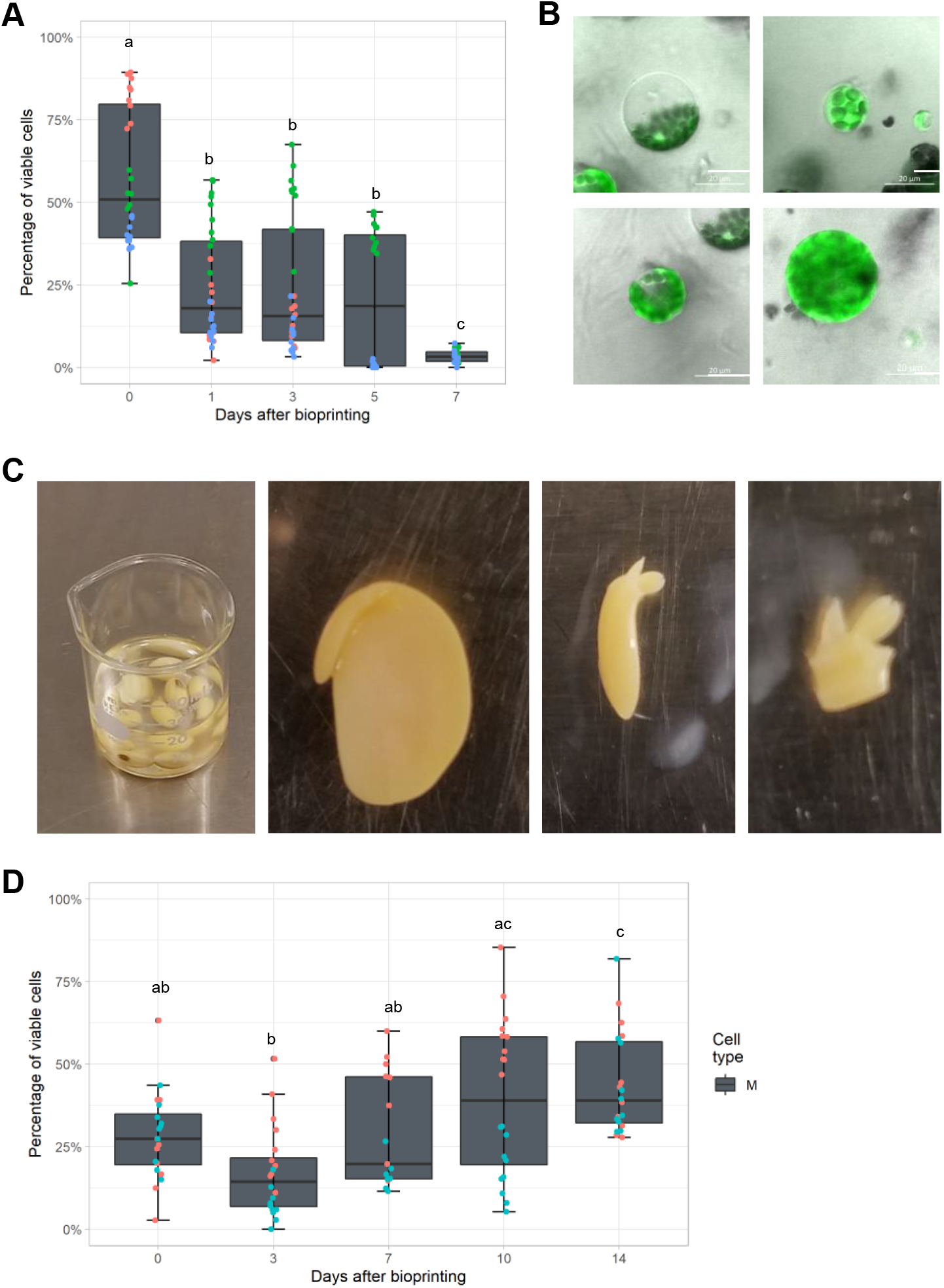
Maintenance of cells isolated from Arabidopsis shoot tissue and soybean meristematic tissue from embryonic axises. (A) Viability of bioprinted shoot-derived bioprinted cells. (B) Representative high resolution confocal images of bioprinted shoot protoplasts. (C) Representative images of soybean tissue dissection (from left to right): mature soybean seeds imbibed in water for 24 hours, dissected imbibed soybean seeds with the embryo exposed, mature soybean embryonic axis, dissected shoot apical meristem and part of hypocotyl with plumules intact. (D) The percentage of viable bioprinted cells from meristematic soybean shoot tissue. Cell viability was evaluated with fluorescein diacetate (FDA) staining immediately after bioprinting (day 0) and 3, 7, 10, and 14 days after bioprinting. Number of bioprinted constructs (N) = 30 (A), N = 20 (D). Letters indicate significant differences (p < 0.05, Tukey test).

**Supplementary Figure S3.**
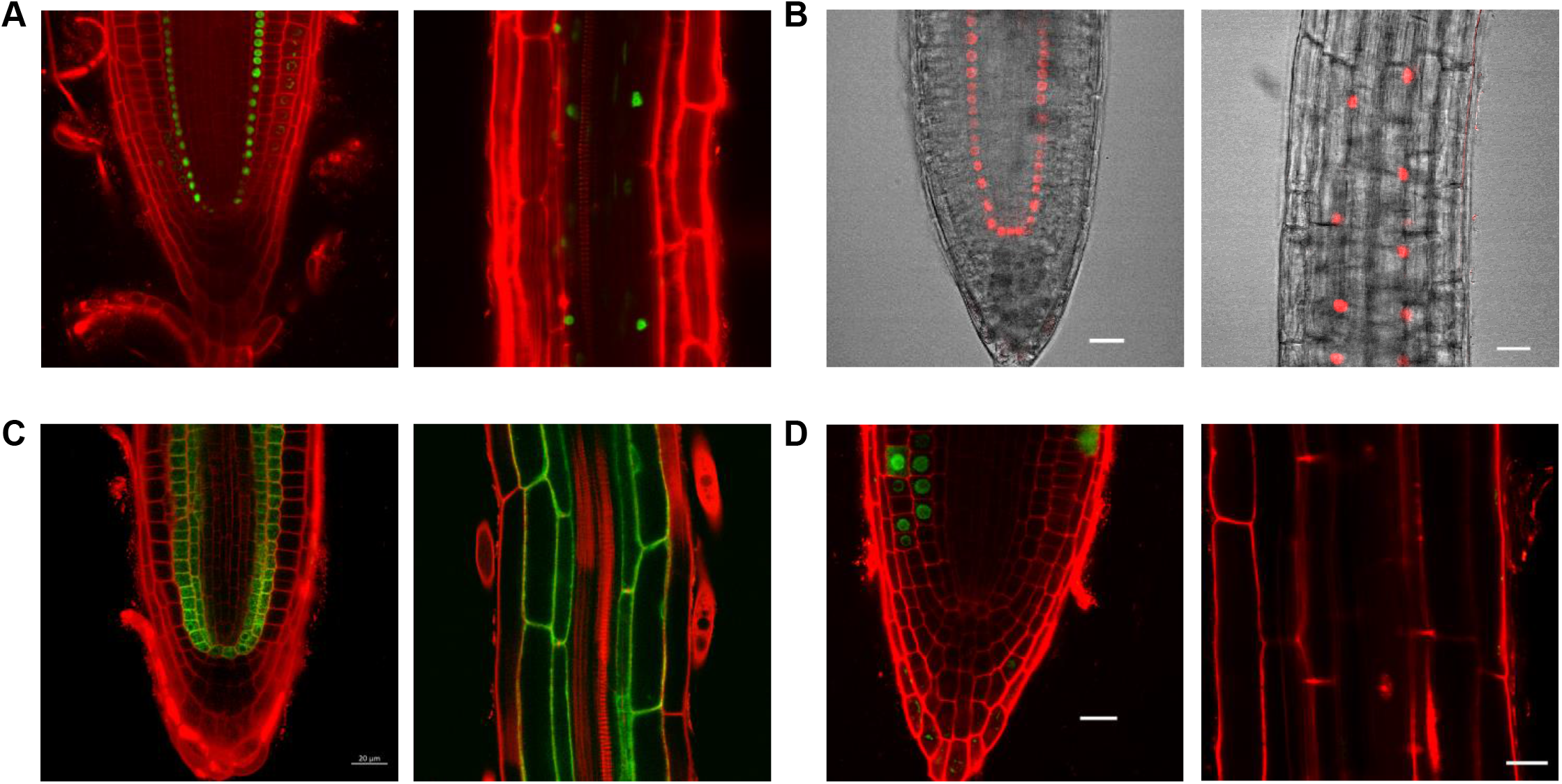
Expression patterns of pEN7:GFP, pSCR:SCR-mCherry, J0571, and pCYCB1;1:CYCB1;1-GFP in the root. Representative confocal images of the meristematic (left) and differentiated (right) region of the root of pEN7:GFP (A), pSCR:SCR-mCherry (B), J0571 (C), and pCYCB1;1:CYCB1;1-GFP (D).

**Supplementary Table S1.** Fractions of endodermal and ground tissue cells in an intact root. The total area, the area of the left and right endodermis cell file, and the area of the left and right ground tissue file in confocal images of a longitudinal section of the root tip were measured in pixels and the ratios to identify the fractions of endodermal and ground tissue cells were calculated.

| Image0 | Total_area | Endodermis_L | Endodermis_R | Ground_L | Ground_R | %Endodermis | %Ground |
| --- | --- | --- | --- | --- | --- | --- | --- |
| 1 | 21613.436 | 1552.741 | 1400.914 | 2275.343 | 2178.491 | 14% | 21% |
| 2 | 20279.388 | 1097.259 | 1465.712 | 1893.104 | 2185.212 | 13% | 20% |
| 3 | 18890.71 | 1392.469 | 1339.218 | 2094.908 | 2032.695 | 14% | 22% |
| 4 | 18504.162 | 1413.839 | 1149.132 | 2040.106 | 1660.279 | 14% | 20% |
| 5 | 19728.087 | 1585.313 | 863.055 | 2482.835 | 1694.229 | 12% | 21% |
| 6 | 21463.849 | 1682.855 | 1767.299 | 2448.713 | 2465.429 | 16% | 23% |
| 7 | 22572.682 | 1345.157 | 879.247 | 2305.795 | 2089.118 | 10% | 19% |
| 8 | 21500.042 | 1834.992 | 1241.306 | 2467.026 | 2280.205 | 14% | 22% |
| 9 | 19330.39 | 1324.965 | 1462.024 | 2008.48 | 2518.796 | 14% | 23% |
| 10 | 16416.475 | 983.435 | 1102.568 | 1564.002 | 1562.365 | 13% | 19% |
|  |  |  |  |  |  | 13% | 21% |

## References

1. M. Ikeuchi, Y. Ogawa, A. Iwase, K. Sugimoto, Plant regeneration: cellular origins and molecular mechanisms. Development. 143, 1442–1451 (2016).

2. K. Duval, H. Grover, L.-H. Han, Y. Mou, A. F. Pegoraro, J. Fredberg, Z. Chen, Modeling Physiological Events in 2D vs. 3D Cell Culture. Physiology. 32, 266–277 (2017).

3. I. Chiesa, C. De Maria, A. Lapomarda, G. M. Fortunato, F. Montemurro, R. Di Gesù, R. S. Tuan, G. Vozzi, R. Gottardi, Endothelial cells support osteogenesis in an in vitro vascularized bone model developed by 3D bioprinting. Biofabrication. 12, 025013 (2020).

4. C. S. Ong, P. Yesantharao, C. Y. Huang, G. Mattson, J. Boktor, T. Fukunishi, H. Zhang, N. Hibino, 3D bioprinting using stem cells. Pediatr. Res. 83, 223–231 (2018).

5. X. Ma, J. Liu, W. Zhu, M. Tang, N. Lawrence, C. Yu, M. Gou, S. Chen, 3D bioprinting of functional tissue models for personalized drug screening and in vitro disease modeling. Adv. Drug Deliv. Rev. 132, 235–251 (2018).

6. E. Maloney, C. Clark, H. Sivakumar, K. Yoo, J. Aleman, S. A. P. Rajan, S. Forsythe, A. Mazzocchi, A. W. Laxton, S. B. Tatter, R. E. Strowd, K. I. Votanopoulos, A. Skardal, Immersion Bioprinting of Tumor Organoids in Multi-Well Plates for Increasing Chemotherapy Screening Throughput. Micromachines (Basel). 11 (2020), doi:10.3390/mi11020208.

7. F. Pampaloni, E. G. Reynaud, E. H. K. Stelzer, The third dimension bridges the gap between cell culture and live tissue. Nat. Rev. Mol. Cell Biol. 8, 839–845 (2007).

8. M. J. Bissell, A. Rizki, I. S. Mian, Tissue architecture: the ultimate regulator of breast epithelial function. Curr. Opin. Cell Biol. 15, 753–762 (2003).

9. J. A. Hickman, R. Graeser, R. de Hoogt, S. Vidic, C. Brito, M. Gutekunst, H. van der Kuip, IMI PREDECT Consortium, Three-dimensional models of cancer for pharmacology and cancer cell biology: capturing tumor complexity in vitro/ex vivo. Biotechnol. J. 9, 1115–1128 (2014).

10. H. Gudapati, M. Dey, I. Ozbolat, A comprehensive review on droplet-based bioprinting: Past, present and future. Biomaterials. 102, 20–42 (2016).

11. T. Pasternak, I. A. Paponov, S. Kondratenko, Optimizing Protocols for Arabidopsis Shoot and Root Protoplast Cultivation. Plants. 10 (2021), doi:10.3390/plants10020375.

12. S. Ochatt, C. Conreux, R. Moussa Mcolo, G. Despierre, J.-B. Magnin-Robert, B. Raffiot, Phytosulfokinealpha, an enhancer of in vitro regeneration competence in recalcitrant legumes. Plant Cell Tissue Organ Cult. 135, 189–201 (2018).

13. M. Benmoussa, S. Mukhopadhyay, Y. Desjardins, Factors influencing regeneration from protoplasts of Asparagus densiflorus cv. Sprengeri. Plant Cell Rep. 17, 123–128 (1997).

14. H. Aoyagi, H. Tanaka, Measurement of viable plant cell and protoplast concentrations with specialized fluorometer. J. Ferment. Bioeng. 77, 517–521 (1994).

15. G. R. López-Marcial, A. Y. Zeng, C. Osuna, J. Dennis, J. M. García, G. D. O’Connell, Agarose-Based Hydrogels as Suitable Bioprinting Materials for Tissue Engineering. ACS Biomater Sci Eng. 4, 3610–3616 (2018).

16. Y. Sakamoto, A. Kawamura, T. Suzuki, S. Segami, M. Maeshima, S. Polyn, L. De Veylder, K. Sugimoto, Transcriptional activation of auxin biosynthesis drives developmental reprogramming of differentiated cells. bioRxiv (2021), p. 2021.06.26.450054.

17. K. Sugimoto, Y. Jiao, E. M. Meyerowitz, Arabidopsis regeneration from multiple tissues occurs via a root development pathway. Dev. Cell. 18, 463–471 (2010).

18. A. Schnittger, L. De Veylder, The Dual Face of Cyclin B1. Trends Plant Sci. 23, 475–478 (2018).

19. M. Xu, Q. Du, C. Tian, Y. Wang, Y. Jiao, Stochastic gene expression drives mesophyll protoplast regeneration. Sci Adv. 7 (2021), doi:10.1126/sciadv.abg8466.

20. C. N. Stewart Jr, M. J. Adang, J. N. All, H. R. Boerma, G. Cardineau, D. Tucker, W. A. Parrott, Genetic transformation, recovery, and characterization of fertile soybean transgenic for a synthetic Bacillus thuringiensis cryIAc gene. Plant Physiol. 112, 121–129 (1996).

21. J. J. Finer, M. D. McMullen, Transformation of soybean via particle bombardment of embryogenic suspension culture tissue. In Vitro Cell. Dev. Biol. Plant. 27, 175–182 (1991).

22. A. Droste, G. Pasquali, M. H. Bodanese-Zanettini, Euphytica. 127, 367–376 (2002).

23. C. M. Hernandez-Garcia, A. P. Martinelli, R. A. Bouchard, J. J. Finer, A soybean (Glycine max) polyubiquitin promoter gives strong constitutive expression in transgenic soybean. Plant Cell Rep. 28, 837–849 (2009).

24. B. Wiebke-Strohm, A. Droste, G. Pasquali, M. B. Osorio, L. Bücker-Neto, L. M. P. Passaglia, M. Bencke, M. S. Homrich, M. Margis-Pinheiro, M. H. Bodanese-Zanettini, Transgenic fertile soybean plants derived from somatic embryos transformed via the combined DNA-free particle bombardment and Agrobacterium system. Euphytica. 177, 343–354 (2011).

25. M. S. Homrich, B. Wiebke-Strohm, R. L. M. Weber, M. H. Bodanese-Zanettini, Soybean genetic transformation: A valuable tool for the functional study of genes and the production of agronomically improved plants. Genet. Mol. Biol. 35, 998–1010 (2012).

26. M. S. Homrich, L. M. P. Passaglia, J. F. Pereira, P. F. Bertagnolli, G. Pasquali, M. A. Zaidi, I. Altosaar, M. H. Bodanese-Zanettini, Resistance to Anticarsia gemmatalis Hübner (Lepidoptera, Noctuidae) in transgenic soybean (Glycine max (L.) Merrill Fabales, Fabaceae) cultivar IAS5 expressing a modified Cry1Ac endotoxin. Genet. Mol. Biol. 31, 522–531 (2008).

27. C. Wu, J. M. Chiera, P. P. Ling, J. J. Finer, Isoxaflutole treatment leads to reversible tissue bleaching and allows for more effective detection of GFP in transgenic soybean tissues. In Vitro Cellular & Developmental Biology - Plant. 44, 540–547 (2008).

28. B. A. McBlain, R. J. Fioritto, S. K. St. Martin, A. J. Calip-Dubois, A. F. Schmitthenner, R. L. Cooper, R. J. Martin, Registration of “Thorne” soybean. Crop Sci. 33, 1406–1406 (1993).

29. W. Lin, J. T. Odell, R. M. Schreiner, Soybean protoplast culture and direct gene uptake and expression by cultured soybean protoplasts. Plant Physiol. 84, 856–861 (1987).

30. J. Haseloff, E. L. Dormand, A. H. Brand, Live imaging with green fluorescent protein. Methods Mol. Biol. 122, 241–259 (1999).

31. Y. Luo, A. Lode, A. R. Akkineni, M. Gelinsky, Concentrated gelatin/alginate composites for fabrication of predesigned scaffolds with a favorable cell response by 3D plotting. RSC Adv. 5, 43480–43488 (2015).

32. S. K. Dhir, S. Dhir, J. M. Widholm, Regeneration of fertile plants from protoplasts of soybean (Glycine max L. Merr.): genotypic differences in culture response. Plant Cell Rep. 11, 285–289 (1992).

33. R. Heidstra, D. Welch, B. Scheres, Mosaic analyses using marked activation and deletion clones dissect Arabidopsis SCARECROW action in asymmetric cell division. Genes Dev. 18, 1964–1969 (2004).

34. N. M. Clark, E. Hinde, C. M. Winter, A. P. Fisher, G. Crosti, I. Blilou, E. Gratton, P. N. Benfey, R. Sozzani, Tracking transcription factor mobility and interaction in Arabidopsis roots with fluorescence correlation spectroscopy. Elife. 5 (2016), doi:10.7554/eLife.14770.

35. E. Buckner, I. Madison, H. Chou, A. Matthiadis, C. E. Melvin, R. Sozzani, C. Williams, T. A. Long, Automated Imaging, Tracking, and Analytics Pipeline for Differentiating Environmental Effects on Root Meristematic Cell Division. Front. Plant Sci. 10, 1487 (2019).

36. J. Park, S. Choi, S. Park, J. Yoon, A. Y. Park, S. Choe, DNA-Free Genome Editing via Ribonucleoprotein (RNP) Delivery of CRISPR/Cas in Lettuce. Methods Mol. Biol. 1917, 337–354 (2019).

37. J. Park, S. Choe, DNA-free genome editing with preassembled CRISPR/Cas9 ribonucleoproteins in plants. Transgenic Res. 28, 61–64 (2019).

40. A. Kiełkowska, A. Adamus, Peptide Growth Factor Phytosulfokine-α Stimulates Cell Divisions and Enhances Regeneration from B. oleracea var. capitata L. Protoplast Culture. J. Plant Growth Regul. 38, 931–944 (2019).

41. C. Rueden, E. Evans, L. Yang, M. Pinkert, Y. Liu, M. Hiner, J. Eglinger, H. Mary, Macarse D. Hereñú, P. Hanslovsky, W. Ouyang, imagej/pyimagej: v1.0.2 (2021; https://zenodo.org/record/5537065).

42. G. Bradski, The openCV library. Dr. Dobb’s Journal: Software Tools for the Professional Programmer. 25, 120–123 (2000).

43. E. Katrukha, ComDet plugin for ImageJ v0.5.3 Zenodo (2020).

44. W. Chang, J. Cheng, J. J. Allaire, C. Sievert, B. Schloerke, Y. Xie, J. Allen, J. McPherson, A. Dipert, B. Borges, shiny: Web Application Framework for R (2021), (available at https://CRAN.R-project.org/package=shiny).

